# A spatial correlation function approach reveals zonation loss in liver disease

**DOI:** 10.64898/2026.11.16.767021

**Authors:** Krishna Chaitanya Ginne, Derek Bowman, Jordi Piñero, Jorge Cobos-Figueroa, Jorge Cañas-Martín, Alba Capelo-Diz, Marta Varela-Rey, Christoph Kuppe, Adrian Ruiz-Villalba, Sudin Bhattacharya, Carlo Piermarocchi

## Abstract

Spatial transcriptomics measures the expression of hundreds to thousands of genes in each cell together with the cell’s position in the tissue. Here we use spatial correlation functions, which describe how the correlation of gene expression between cells decays with distance, to quantify how the spatial organization of gene expression changes in disease. From the correlation function of each gene, or of each pair of genes, we extract two numbers: a susceptibility, which measures how strongly gene expression is correlated between nearby cells, and a correlation length, the distance over which this correlation extends. Both are in physical units and can be compared directly between samples and conditions. We apply the approach to healthy and fibrotic mouse liver using newly generated data, and to healthy and fibrotic human liver using published data. Correlations between gene pairs identify genes of the same hepatocyte zone, and assign a zone to 1,266 genes that are not known zonation markers. The two fibrotic conditions differ: CCl_4_ injection in mice produces a moderate reorganization in which some genes lose spatial correlation and others gain it, while in human fibrosis almost every measured gene loses it. Changes in spatial correlation are only weakly related to changes in expression level. For instance, *Cyp2c29* and *Pon1* partially lose their spatial correlation in treated mice, although their average expression does not consistently fall. Spatial correlation functions therefore detect changes in tissue organization that measurements of expression level alone miss.

## Introduction

Complex biological tissues can be seen as the living counterparts of complex heterogeneous materials (Torquato 2002). In both tissues and materials, many heterogeneous components interact collectively and give rise to emergent properties that define the global behavior of the system. In complex materials, it is not enough to look at the presence of domains or interfaces; one also needs to examine how spatial fluctuations of a variable of interest around those average patterns are correlated. This introduces the concept of a correlation length, which defines a characteristic spatial range over which fluctuations are correlated. The correlation length indicates the degree of cooperation among spatially varying quantities across scales, helps to distinguish ordered and disordered phases, and is often enhanced near transition points (Solé 2011). Correlated fluctuations can grow even before clear structures or domains emerge, revealing early signs of collective behavior and the system’s proximity to an organized state. A similar perspective can be brought to biology. Spatial transcriptomics at single-cell resolution provides direct access to the spatial organization of gene expression across tissues (Rao et al. 2021), enabling the computation of correlation functions between cells. This makes it possible to use frameworks based on correlation lengths to quantify cellular heterogeneity, identify emergent tissue-scale organization, and detect transitions between physiological and pathological states.

Many computational tools now describe spatial organization in spatial transcriptomics data, from neighborhood and co-occurrence analyses (Dries et al. 2021; Palla et al. 2022) to tests for spatially variable genes (Svensson et al. 2018; Edsgärd et al. 2018; Sun et al. 2020; Zhu et al. 2021) and spatial autocorrelation statistics from the geographical sciences (Zormpas et al. 2023). They are designed mainly to test whether a pattern is present, and they usually summarize it at one chosen scale. Our main object of interest, however, is the gene expression correlation fluctuations as a function of distance. By focusing on how this correlation decays with distance, we treat the biological tissue as a complex material system, enabling the characterization of collective behavior and emergent organization. This is also different from differential expression, which is still the standard way to compare tissue states across conditions. Differential expression measures how much a gene is expressed, but not how its expression is ordered spatially in the tissue: a gene can lose its spatial organization while its abundance does not change. This difference matters biologically, because the function of a tissue depends not only on how much each gene is expressed but on where, and disease can disrupt this spatial organization. In addition, the correlation between pairs of genes reveals which genes are co-expressed in the same region and which ones are mutually exclusive. Therefore, correlation functions can be used to study patterns of genetic correlation along spatial axes, which is particularly interesting for liver zonation.

These quantities have precise definitions in the statistical physics of fluctuating fields (Chaikin and Lubensky 1995; Goldenfeld 1992). For fluctuations *δϕ*(**r**) of a field about its mean, the two-point correlation function *C*(*r*) *∝ ⟨δϕ*(**0**) *δϕ*(**r**)*⟩* measures how strongly a fluctuation at one point is statistically related to the fluctuation at a second point separated by a distance *r*. We divide the two-point averages by the variance of expression, so that the correlation function equals one at zero distance, where a cell is compared with itself, and decreases as the two cells become unrelated. The rate of this decay defines the correlation length *ξ*, the characteristic distance over which fluctuations remain correlated, commonly obtained from an exponential (Ornstein–Zernike (Chaikin and Lubensky 1995)) form *C*(*r*) *∼ e*^*−r/ξ*^. The spatial integral of the correlation function defines the susceptibility *χ* = ∫*C*(*r*) *d*^2^*r*, which accumulates the correlated fluctuations across multiple scales. An analogous approach has been used to characterize collective behavior in other biological systems, in particular in the analysis of velocity fluctuations in bird flocks (Cavagna et al. 2010; Cavagna and Giardina 2014), where the same correlation function and correlation length have been used to study the proximity to a scale-free regime, in which the correlation length diverges.

Here we bring these quantities to spatial transcriptomics, where they can be used not only to detect spatial patterns but also to compare the organization of a tissue between conditions. We build the approach on two correlation functions, which capture different aspects of tissue organization. The single-gene autocorrelation *C*_*g*_(*r*) describes how the expression of one gene is correlated between cells at increasing separation, and identifies genes changing in spatial organization with the biological state. The gene-pair cross-correlation 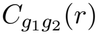 extends this concept to the coordinated expression of two different genes, quantifying the spatial range over which their co-expression or mutual exclusion is maintained. Unlike the single-gene auto-correlation function, this function can detect both the weakening of positive and negative coupling among two genes. From each correlation function, we extract the two quantities introduced above: the correlation length *ξ* gives the characteristic distance over which organization persists and is the more intuitive quantity, identifying contraction or expansion of spatial domains. The susceptibility *χ*, on the other hand, measures the total amount of correlation accumulated up to a fixed range and requires no assumption about the functional form of the decay. We therefore use *ξ* to characterize the domains and rank spatially correlated genes and gene pairs using *χ*, which is robust to highly variable data that is otherwise difficult to fit.

We focus here on the liver, an organ with a structured and well-defined spatial organization in healthy conditions (Halpern et al. 2017). The liver’s geometry consists of hexagonally packed lobules centered around a central vein. A peri-portal to peri-central axis aligns hepatocytes, vascular, and bile networks in a coordinated and zonated pattern. In particular, fibrotic tissue is well suited to a correlation-function framework. This disease induces changes in cellular composition and organization, affecting how signals propagate, and how cells interact with other cells and the matrix (Ramachandran et al. 2019; Kisseleva and Brenner 2021). We apply the correlation function framework to new Xenium data of mouse liver with CCl_4_-induced chronic injury and to published MERFISH data from healthy and fibrotic human liver tissues (Watson et al. 2025). We find that changes in the spatial correlation of gene expression are only weakly related to changes in the level of expression of these genes, and that the sign of gene-pair correlations recovers the zonation of hepatocytes and assigns unannotated genes along the zonation axis. The two fibrotic conditions differ: chronic fibrosis induced by CCl_4_ in mice reorganizes the spatial correlation of hepatocyte genes, while chronic and progressive human fibrosis causes an almost uniform loss.

## Results

### Spatial correlation functions and derived quantities

We define our two-point statistics starting with the single-cell transcriptomic data *X ∈* ℝ^*N ×G*^ (*N* cells and *G* genes) with spatial coordinates **r**_*i*_ *∈* ℝ^2^ for cell *i*. Let *D*_*p*_ denote the set of cell pairs (*i, j*) with distance *s*_*ij*_ = *∥***r**_*i*_ − **r**_*j*_*∥* = *s*_*p*_ in the *p*-th distance bin. We consider two two-point statistics: (1) the gene-centric autocorrelation: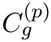, and (2) the gene-pair correlation: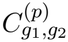.

Let *X*_*i*_(*g*) denote the expression of gene *g* in cell *i*. We center the expression levels as

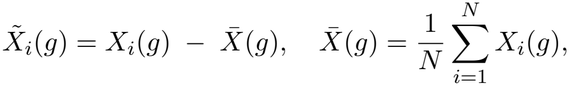

and denote by 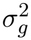 the variance of gene *g* over the *N* cells of the sample. The distance-resolved gene-centric autocorrelation for gene *g* in bin *p* is

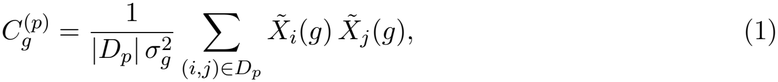

where *D*_*p*_ = {(*i, j*) : *s*_*ij*_ *∈* [*s*_*p−*1_, *s*_*p*_)} is the set of cell pairs whose spatial separation *s*_*ij*_ = *∥***r**_*i*_ − **r**_*j*_*∥* falls in distance bin *p*, and |*D*_*p*_| is the number of such pairs. This function measures how the expression fluctuations of gene *g* remain correlated as a function of spatial separation. The corresponding gene-centric correlation length *ξ*_*g*_ is obtained by fitting the decay of 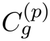 with distance, e.g., via an exponential model

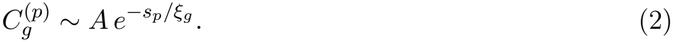

The susceptibility *χ*_*g*_ is the integral of the correlation function over the plane. Writing it as a discrete sum over the bins of distance *s*_*p*_ and width Δ*s*, the two-dimensional radial measure gives

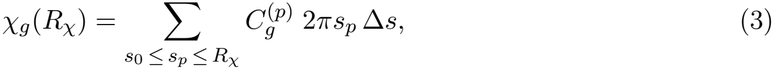

where the sum runs from a short-distance cutoff *s*_0_, set to about the diameter of a human hepatocyte (Rohr et al. 1976) (30 *µ*m, or about two mouse hepatocyte diameters) so that the sum leaves out adjacent cells, up to an upper cutoff *R*_*χ*_. The cutoff *R*_*χ*_ sets the size of the neighborhood over which correlated fluctuations are added up, so that *χ*_*g*_(*R*_*χ*_) measures how coordinated the expression of a gene is between a cell and the cells around it within this distance. We take *R*_*χ*_ equivalent to several hepatocyte diameters. Note that the upper cutoff is part of the definition. The function 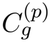 obeys a sum rule that fixes its integral over the entire system independently of the underlying biology, so the unrestricted sum is a constant that carries no information about the gene and becomes negative at large separation. The susceptibility is therefore meaningful only as a partial integral up to a given *R*_*χ*_, and every susceptibility in this work is defined according to the same cutoff. The sum rule, the values of *s*_0_ and *R*_*χ*_ used in our approach, and their justification are given in Methods.

For a gene pair (*g*_1_, *g*_2_), the distance-resolved gene-pair correlation in bin *p* is

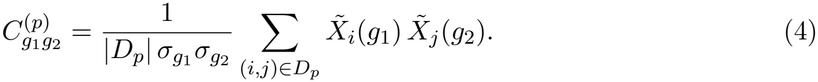

This quantity characterizes the spatial range over which the expression of gene *g*_1_ in one cell is correlated with the expression of gene *g*_2_ in another cell at distance *s*_*p*_. The pair-level correlation length *ξg*_1_*g*_2_ is obtained by fitting the decay of 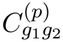 with distance, using the same exponential framework as for the single gene case, and the pair level susceptibility 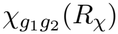 is defined by Eq (3) with 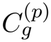 replaced by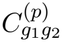, with the same cutoffs. Unlike 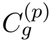, the cross-correlation is not constrained to be positive at short range. Indeed, 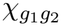 is negative if the sign of fluctuations of the first gene is opposite to the sign of fluctuations of the second gene.

### Single-gene correlation functions in hepatocytes

We start with the correlation functions of individual genes over hepatocytes. For each sample, we calculate the distance-resolved autocorrelation 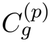of Eq (1) for hepatocytes only, using 5 *µ*m bins out to 300 *µ*m, and extract the correlation length *ξ*_*g*_ by fitting the exponential decay on the interval 30 ≤ *s*_*p*_ ≤ 100 *µ*m. This is the same interval used for the susceptibility (Methods), so that *ξ*_*g*_ and *χ*_*g*_ are estimated consistently.

Figure 1 shows two genes in the Xenium dataset representative of two different qualitative behaviors detected by our analysis. For *Cyp2a5* (Fig. 1A,B) the correlation length contracts in the fibrotic tissue, from *ξ*_*g*_ = 100 ± 23 *µ*m in control to 72 ± 9 *µ*m after CCl_4_ treatment, a ratio of 1.39. For *Etnppl* (Fig. 1C,D), however, the rate of decay does not change: the curves of the two conditions are parallel on the semi-logarithmic scale across the entire fit range, *ξ*_*g*_ = 118 ± 29 *µ*m versus 117 ± 12 *µ*m, a ratio of 1.00, and the values of the individual mice overlap (126 and 111 *µ*m in control, 122 and 108 *µ*m after CCl_4_). Both genes are well described by the exponential model (*R*^2^ *>* 0.94 in all four cases). Figure 1E extends the comparison to the twelve genes with *R*^2^ *>* 0.93 in both conditions, and shows that both behaviors are well represented: most genes contract, while a subset is unchanged or slightly expanded.

**Figure 1.**
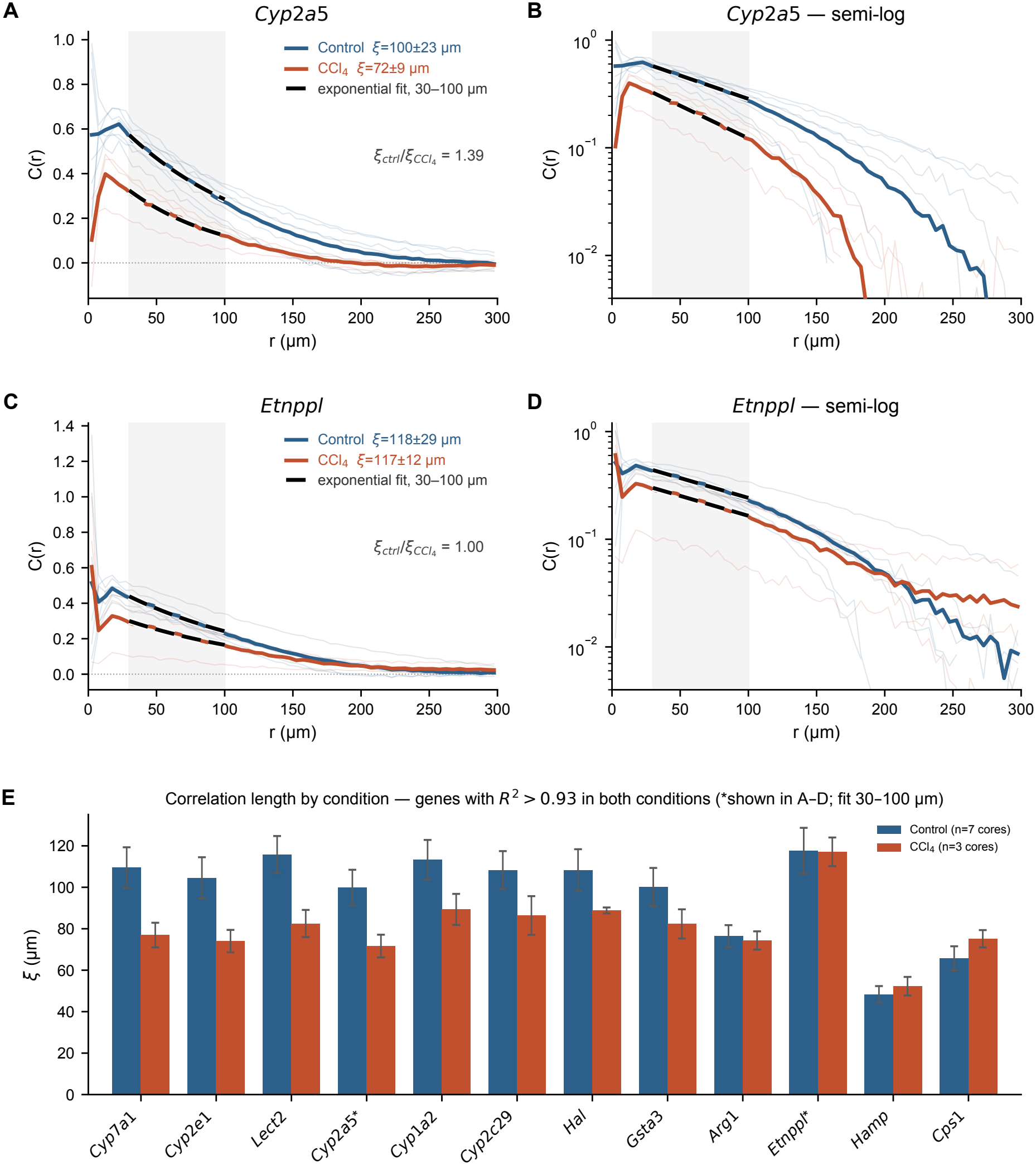
Single-gene spatial correlation functions and correlation lengths in the Xenium liver fibrosis dataset. Two genes are shown as illustrative examples: a gene whose correlation length changes between conditions (*Cyp2a5*) and one whose correlation length does not (*Etnppl*). (A, B) Single-gene 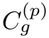 of *Cyp2a5* as a function of pair separation *r*, shown on linear (A) and semi-logarithmic (B) scales. Thin lines are individual tissue cores, bold lines are the mean over cores within each condition (blue, control, *n* = 7 cores from two mice; red, CCl_4_, *n* = 3 cores from two mice), and dashed black lines show the exponential decay evaluated at the mean per-core *ξ* (with amplitude fitted to the condition mean). The shaded band marks the 30–100 *µ*m fit window. Correlation lengths in the legend are the mean ± standard deviation of the per-core fits. (C, D) The same quantities for *Etnppl*, for which the control and CCl_4_ curves decay at the same rate over the fit range. (E) Correlation length by condition for twelve genes having exponential fits with *R*^2^ *>* 0.93 in both conditions, ordered by relative change; error bars are the standard error of the mean across cores, and the asterisks identify the two genes shown in A–D. Both contraction and absence of change are observed within this set.

The correlation length we introduce here is a summary statistic, and it is important to explore how a contraction of *ξ* corresponds to changes in the tissue itself. Figure 2 shows the spatial distribution of expression of the same two genes, in one representative control core and one representative CCl_4_-treated core. These representative cores were chosen as the ones where the fitted *ξ* was closest to the mean value of their respective conditions. The panels show the per-cell expression directly, with the expression standardized to zero mean and unit variance within each core, so that it reflects the spatial pattern of fluctuations rather than differences in overall abundance. The top row shows the same two sections stained with H&E and aligned to the expression panels (Fig. 2A,B).

**Figure 2.**
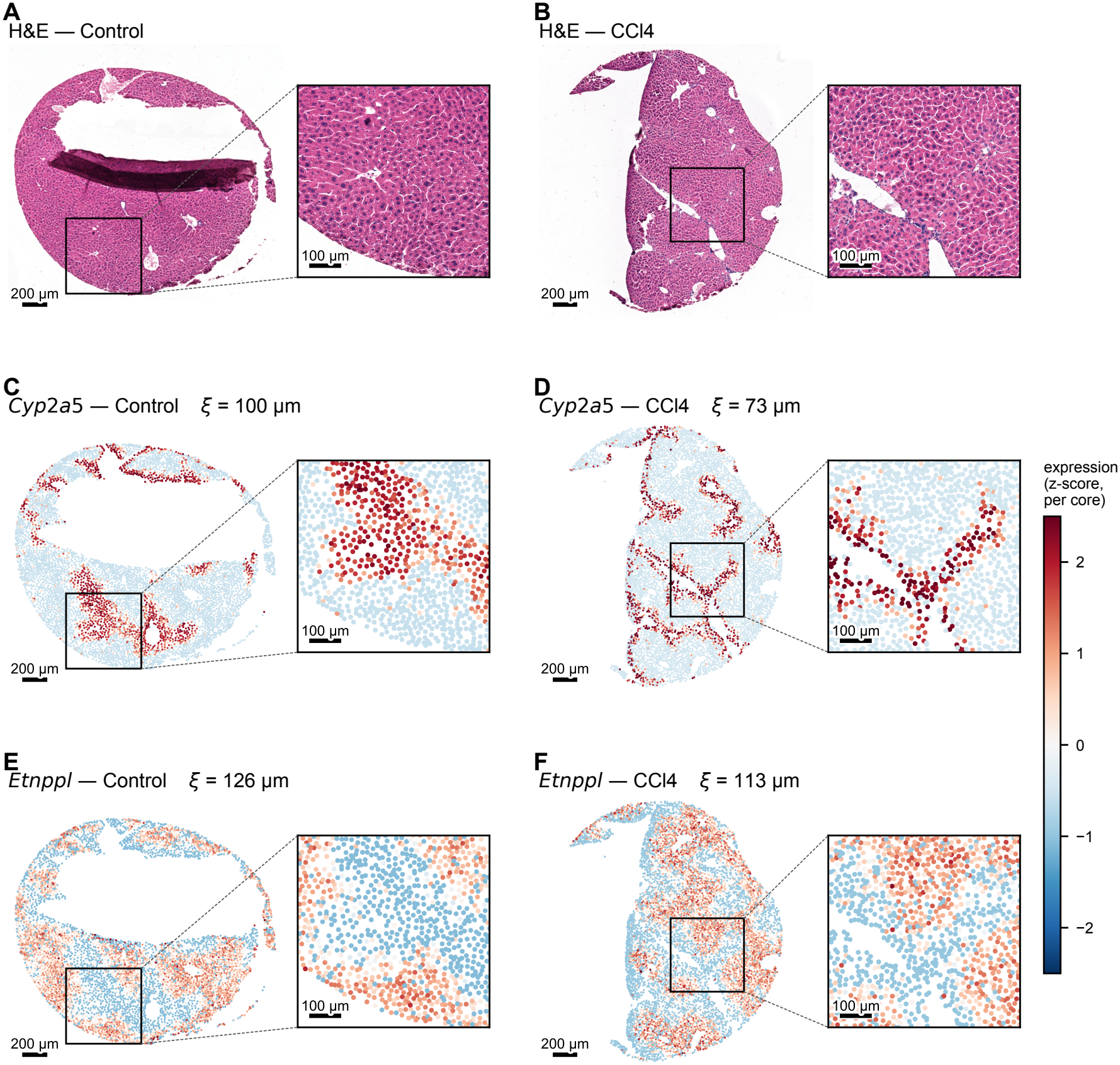
Histology and spatial expression patterns underlying a change in correlation length. One representative control core (`core_0`) and one representative CCl_4_ core (`core_18`). Each panel shows the whole core, with a boxed 600 × 600 *µ*m region magnified on its right; the boxed region is the same in the three rows of each condition. Scale bars, 200 *µ*m (whole core) and 100 *µ*m (magnified region). (A, B) Hematoxylin and eosin (H&E) staining of the same sections, shown as a reference for the tissue architecture. The dark band in the control section (A) is a fold of the tissue formed during staining. (C–F) Per-cell expression of hepatocytes, standardized within each core, with one color scale for all panels. (C, D) *Cyp2a5*, whose correlation length contracts between conditions, forms broad contiguous regions in control tissue (C) and narrow band-like structures after CCl_4_ treatment (D). (E, F) *Etnppl*, whose correlation length is unchanged, looks comparable in both conditions.

For *Cyp2a5* the difference in the pattern of expression between control and CCl_4_ tissue is clear (Fig. 2C,D). In the control tissue the gene is expressed in the hepatic parenchyma and organized in broad, continuous regions of high expression across many tens of microns. After CCl_4_ treatment, the same gene is confined to a network of narrow, band-like structures. On the other hand, *Etnppl*, for which *ξ* is unchanged, is expressed in the hepatic parenchyma in both conditions, without a clear difference in the spatial distribution of its expression (Fig. 2E,F). The reorganization for *Cyp2a5* is not a simple isotropic contraction of otherwise similar domains: the pattern changes character, from compact territories to thin, branching bands.

The individual cores in Fig. 2 were chosen as representative of their condition, but the underlying effect is more variable. Across all cores (Supplemental Fig. S1), the filament-like *Cyp2a5* pattern is reproduced in all three CCl_4_ cores. However, the control cores span a much wider range (72–137 *µ*m) and the lowest-*ξ* core in the control can resemble the fibrotic tissue. The change on injury is therefore a shift in the distribution of *ξ* for this gene, with partial overlap between conditions (Supplemental Fig. S1B), rather than a systematic switch.

### CCl_4_ injury reorganizes spatial correlation without a net loss in expression

For every gene that passes the fit-quality filters, we compare the susceptibility between control and CCl_4_ tissue. We rank genes by *Z*_*χ*_, a z-score that divides the change in *χ* between the two conditions by its variability between tissue cores (Methods), so that a gene ranks highly when the change is large compared with this variability, and a positive value indicates a loss of spatial correlation. As a summary of how spatial organization changes after injury (Fig. 3A), we count the genes showing a *positive-correlation loss* (i.e. the genes above the diagonal line), for which 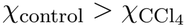, indicating a reduction in spatial correlation in fibrotic tissue compared to control. Of the 297 genes passing the filters, 147 (49%) show a positive-correlation loss, with a median ratio of 1.42; the remainder are unchanged or gain correlation, and points fall on both sides of the diagonal (Fig. 3A). This loss is not spread at random across genes: it includes every known zonation marker on the panel. We use a curated set of sixteen confident mouse hepatocyte zonation markers, selected from the continuous porto-central expression profiles of published spatial studies of the mouse liver (Halpern et al. 2017; Hildebrandt et al. 2021; Amberg et al. 2026). These include the periportal markers *Sds, Ass1, Asl, Pck1, Cyp2f2, Hal, Alb, Arg1, Hsd17b13* and *Cps1*, and the pericentral markers *Glul, Cyp2e1, Cyp1a2, Oat, Slc1a2* and *Cyp2a5*. Six of these are represented on the 5,046-gene Xenium panel: the periportal markers *Arg1, Cps1* and *Hal*, and the pericentral markers *Cyp1a2, Cyp2a5* and *Cyp2e1*. All six are among the 147 genes that show a positive-correlation loss, with a median ratio 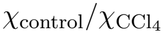 of 1.56, against 1.42 for the 147 genes as a whole: the pericentral markers *Cyp2a5* (2.03×), *Cyp1a2* (1.85×) and *Cyp2e1* (1.33×), and the periportal markers *Cps1* (1.75×), *Arg1* (1.36×) and *Hal* (1.27×). Apart from *Cyp1a2*, the highest-ranking genes overall (Table 1 and Supplemental Fig. S2) are not zonation markers. They include the MAPK/ERK genes *Egr1* and *Dusp6*, metabolic genes such as *Cyp2c29, Cyb5a* and *Slco1a1*, and the hepatocyte-secreted *Fgl1*.

**Table 1.** Mouse CCL_4_ dataset: genes with the largest change in spatial correlation after CCl_4_. The ten highest-ranking genes by *Z*_*χ*_. *χ* is the susceptibility (Eq 3) evaluated over 30 ≤ *r* ≤ 100 *µ*m and averaged over the tissue cores of each condition; the ratio is 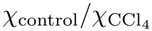, so values *>* 1 indicate a loss of spatial correlation in disease. *Z*_*χ*_ (Eq 5) normalizes the difference by the between-core variability. log_2_FC is the differential-expression fold-change for the same cells, positive when the gene is more highly expressed in control tissue, and “DE %ile” is the gene’s percentile within the fold-change ranking (0% = largest fold-change); a high DE percentile therefore identifies a gene that reorganizes spatially while its mean abundance barely changes. “n.d.” marks genes whose *χ* is not positive in both conditions, for which a ratio is not meaningful. Across the whole panel, 147 of 297 genes show a positive-correlation loss (median 1.42×). Complete statistics for all genes are given in Supplemental Table S1.

| Gene | Zone | χ control | χ CCl<sub>4</sub> | ratio | *Z*<sub>χ</sub> | log<sub>2</sub>FC | DE %ile |
|---|---|---|---|---|---|---|---|
| *Cyb5a* | — | 2,609 | 1,582 | 1.6 | 2.7 | 0.96 | 19 |
| *Cyp2c29* | — | 13,098 | 6,743 | 1.9 | 2.4 | 0.28 | 96 |
| *Egr1* | — | 1,042 | 67 | 15.6 | 2.4 | 2.05 | 1 |
| *Sntb1* | — | 1,525 | 448 | 3.4 | 2.4 | 0.50 | 79 |
| *Plpp3* | — | 1,994 | 651 | 3.1 | 2.4 | 1.03 | 14 |
| *Fgl1* | — | 1,136 | 555 | 2.0 | 2.4 | 1.50 | 3 |
| *Cyp1a2* | Central | 12,069 | 6,511 | 1.9 | 2.1 | 0.50 | 80 |
| *Slco1a1* | — | 1,510 | 526 | 2.9 | 2.1 | 1.41 | 4 |
| *Dusp6* | — | 1,280 | 395 | 3.2 | 2.1 | 1.51 | 3 |
| *Nrn1* | — | 4,338 | 2,430 | 1.8 | 2.0 | 0.47 | 83 |

**Figure 3.**
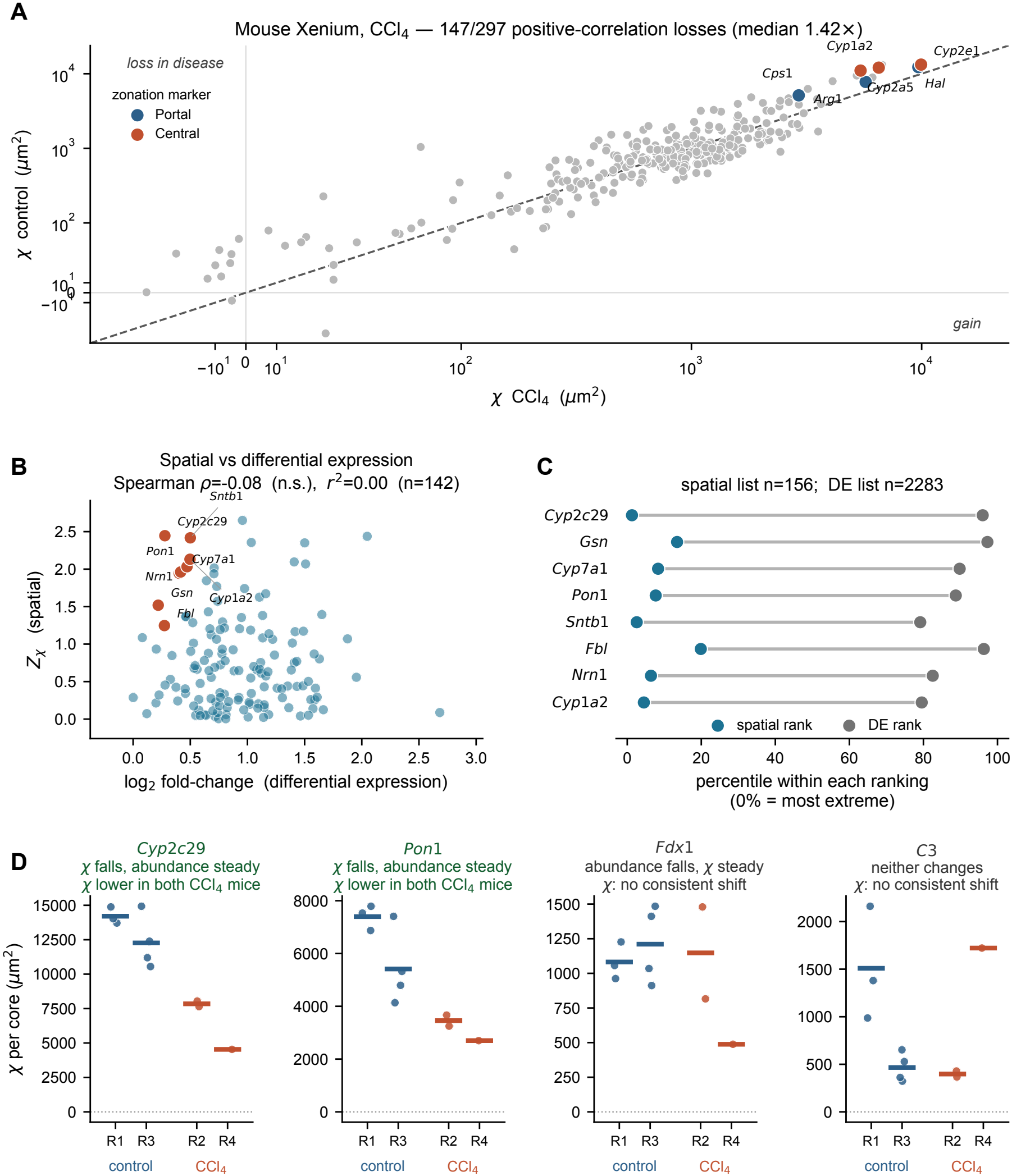
CCl_4_ injury reorganizes spatial correlation without a net loss. (A) Susceptibility *χ* per gene (Eq 3), averaged over tissue cores. Genes above the diagonal lost spatial correlation after injury. Axes are on a symmetric-log scale, linear within ±25 *µ*m^2^ of zero, so the few genes with small negative *χ* are displayed. 147 of 297 genes show a positive-correlation loss (median 1.42×). Curated mouse zonation markers are colored by zone and all six lose correlation. (B) *Z*_*χ*_ against log_2_ fold-change. The two statistics show no rank association among the genes with *Z*_*χ*_ *>* 0 and a positive log_2_ fold-change; the eight genes with the largest normalized rank difference are highlighted and labeled. (C) Position of those eight genes within each ranking, as a percentile of the corresponding list (0% = most extreme). A long connecting bar marks a gene the spatial analysis prioritizes and a fold-change analysis does not. (D) *χ* in each tissue core (points) and its mean for each mouse (bars) for four genes that illustrate the dissociation between spatial correlation and abundance; R1 and R3 are the control mice, R2 and R4 the CCl_4_ mice (R4 has low transcript detection). *Cyp2c29* and *Pon1* have lower *χ* in both CCl_4_ mice although their abundance does not fall consistently; *Fdx1*, whose abundance does fall, and *C3* show no consistent shift in *χ*. Note the independent vertical scales.

This loss of spatial correlation is weakly related to differential expression (DE). Among the 142 genes with *Z*_*χ*_ *>* 0 and a positive log_2_ fold-change between all control and all CCl_4_ hepatocytes, pooled across tissue cores, the two rankings show no association (*ρ* = −0.08, *p* = 0.36; *r*^2^ = 0.00; Fig. 3B). Some genes at the very top of the spatial ranking sit near the bottom of the DE ranking: *Cyp2c29* is 2nd of 156 by *Z*_*χ*_ but 2192nd of 2283 by fold-change, and *Gsn, Cyp7a1* and *Pon1* behave similarly (Fig. 3C). Fourteen of the 156 genes in the spatial list do not appear in the fold-change list at all. These genes reorganize spatially while their mean expression over hepatocytes changes little, so a ranking by fold-change would not bring them forward.

Averaging the cores of each mouse also allows the distinction from differential expression to be checked mouse by mouse (Fig. 3D). *Cyp2c29* and *Pon1* lose spatial correlation in both CCl_4_ mice: their mean *χ* is 7,846 and 4,537 *µ*m^2^ (*Cyp2c29*) and 3,457 and 2,700 *µ*m^2^ (*Pon1*), against 14,207 and 12,267 *µ*m^2^, and 7,398 and 5,415 *µ*m^2^, in the two control mice. Their mean expression, in contrast, does not fall consistently across the two CCl_4_ mice, and the fall seen when cells are pooled comes largely from R4, the mouse with low transcript detection. *Fdx1* shows the opposite dissociation: its expression falls in both CCl_4_ mice, while its *χ* is unchanged in R2 and lower only in R4. *C3* changes in neither respect.

### Gene-pair correlations recover the zonation axis and place unannotated genes along it

The cross-correlation 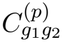 of Eq. (4) measures how the expression of one gene in one cell is related to that of a second gene in another cell at a distance *s*_*p*_. Unlike the single-gene function it is signed. Cells separated by less than the porto-central distance tend to lie in a similar zone, so over the range of *χ* two genes with the same porto-central gradient have 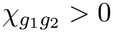, whereas two genes with opposite gradients have 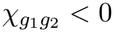. This applies only to strongly coupled pairs, because most pairs join genes with weak porto-central gradients and have a small positive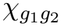. Loss of coupling moves 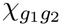 toward zero in either case, from above for same gradients and from below for opposite gradients.

We evaluated all 251,244 gene pairs in the mouse panel and partitioned them into two disjoint sets on this basis. The *positive-coupling loss* set requires *χ >* 0 in both conditions, and contains 72,742 pairs. The *negative-coupling loss* set requires *χ <* 0 in both conditions, and contains 11,222 pairs. Six of the curated zonation markers introduced above are on the panel: three periportal (*Arg1, Cps1, Hal*) and three pericentral (*Cyp1a2, Cyp2a5, Cyp2e1*). Together they form 15 pairs whose expected sign is known in advance: the 6 pairs of markers from the same zone (three in each zone) should be co-localized, and the 9 pairs of markers from opposite zones should be anti-localized. All 15 pairs behave as expected, with the 6 same-zone pairs in the positive set and the 9 opposite-zone pairs in the negative set (Fig. 4B, Table 2). The sign is the same in each of the ten tissue cores, so this relationship does not depend on variability between cores or mice. The sign of the cross-correlation therefore recovers the zonation relationship. Specifically, a positive cross-correlation indicates that the genes are within the same zone, while a negative cross-correlation indicates that the genes are in different zones.

**Table 2.** Gene pairs among the mouse hepatocyte zonation markers. All 15 pairs formed between confident zonation markers that pass either coupling filter, from the 251,244 gene pairs evaluated in the mouse Xenium cohort. Pairs are assigned to the positive-coupling set (*χ >* 0 in both conditions and *Z*_*χ*_ *>* 0; 72,742 pairs in total) or the negative-coupling set (*χ <* 0 in both conditions and *Z*_*χ*_ *<* 0; 11,222 pairs). The ratio is 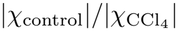, so values *>* 1 indicate a weakening of the coupling, whatever its sign. Every pair in the positive set consists of two markers of the same zone and every pair in the negative set consists of markers of opposite zones, with no exceptions in either direction.

| Pair | Zones | χ control | χ CCl<sub>4</sub> | \|χ\| ratio | *Z*<sub>χ</sub> |
|---|---|---|---|---|---|
| **Negative coupling loss** — mutually excluded pairs, all opposite-zone |  |  |  |  |  |
| *Cyp1a2–Hal* | Central–Portal | -11,794 | -7,333 | 1.61 | -1.48 |
| *Cyp2a5–Hal* | Central–Portal | -10,973 | -6,773 | 1.62 | -1.27 |
| *Arg1–Cyp2a5* | Central–Portal | -8,763 | -5,156 | 1.70 | -1.27 |
| *Arg1–Cyp1a2* | Central–Portal | -8,144 | -5,385 | 1.51 | -1.25 |
| *Cps1–Cyp2a5* | Central–Portal | -6,587 | -3,341 | 1.97 | -1.18 |
| *Cps1–Cyp1a2* | Central–Portal | -5,698 | -3,380 | 1.69 | -1.09 |
| *Cyp2e1–Hal* | Central–Portal | -12,224 | -9,128 | 1.34 | -0.88 |
| *Arg1–Cyp2e1* | Central–Portal | -7,777 | -6,149 | 1.26 | -0.69 |
| *Cps1–Cyp2e1* | Central–Portal | -5,213 | -3,597 | 1.45 | -0.66 |
| **Positive coupling loss** — co-localized pairs, all same-zone |  |  |  |  |  |
| *Arg1–Hal* | Portal–Portal | 8,595 | 6,985 | 1.23 | +0.60 |
| *Cps1–Hal* | Portal–Portal | 6,275 | 4,650 | 1.35 | +0.63 |
| *Arg1–Cps1* | Portal–Portal | 6,079 | 3,838 | 1.58 | +0.91 |
| *Cyp2a5–Cyp2e1* | Central–Central | 10,680 | 6,606 | 1.62 | +1.57 |
| *Cyp1a2–Cyp2e1* | Central–Central | 12,328 | 7,598 | 1.62 | +1.76 |
| *Cyp1a2–Cyp2a5* | Central–Central | 10,896 | 5,727 | 1.90 | +2.03 |

**Figure 4.**
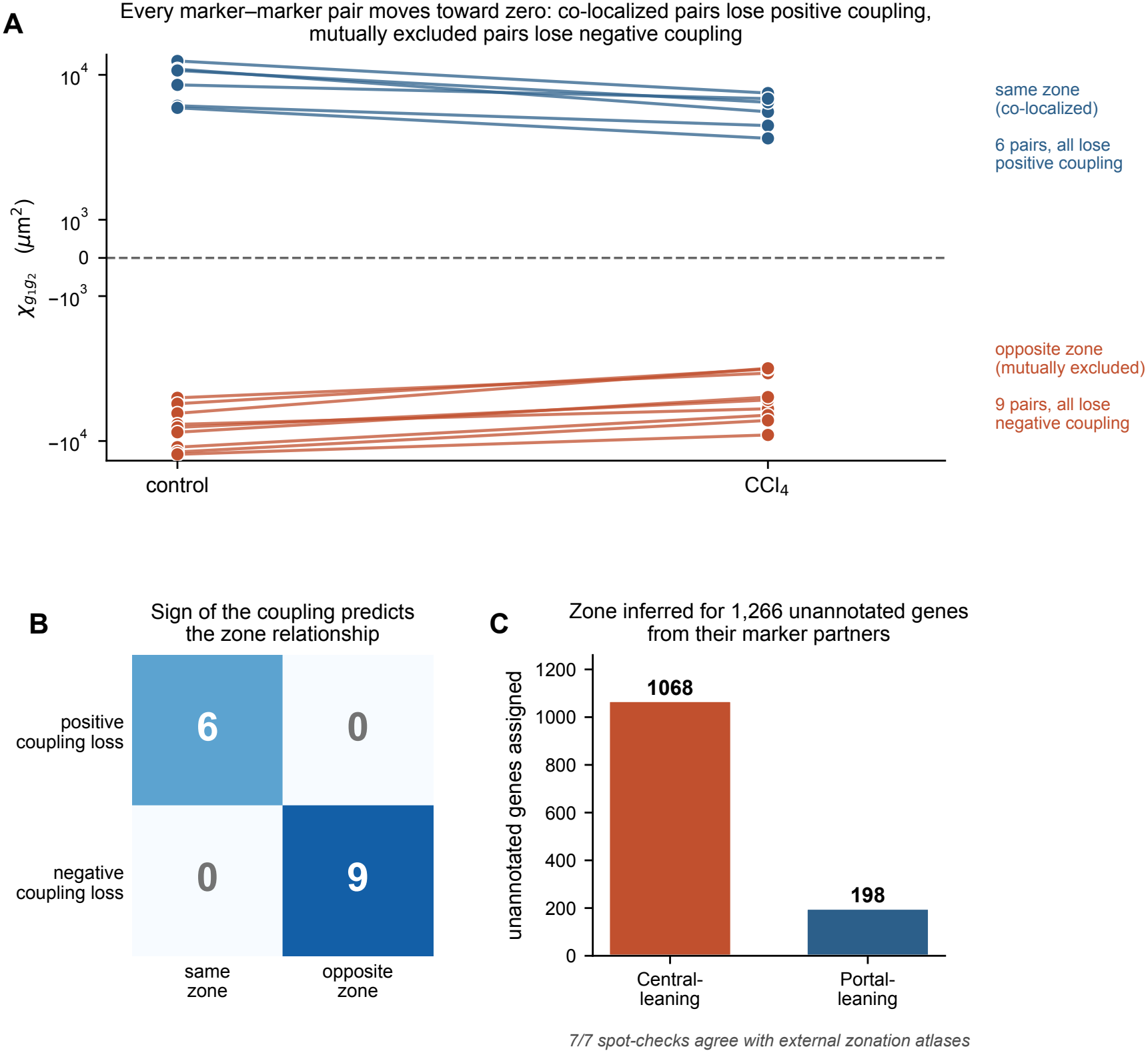
Gene-pair correlations recover the zonation axis and its degradation after CCl_4_ injury. (A) Pair susceptibility 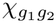 in control and after CCl_4_ for the 15 pairs formed between confident mouse zonation markers, on a symmetric-log axis (linear within ×2 ± 10^3^ *µ*m^2^) so that co-localized and anti-localized pairs are shown together. Every pair moves toward zero: positive couplings weaken from above, negative couplings from below. (B) The sign of the coupling predicts the zone relationship. Of the marker–marker pairs passing the positive-coupling filter, all six join markers of the same zone and none join opposite zones; of those passing the negative-coupling filter, all nine join opposite zones and none the same zone. (C) Zone assigned to 1,266 unannotated genes through their marker partners, taking co-localization as evidence for the marker’s zone and anti-localization as evidence for the opposite zone.

Both halves of the axis degrade together. Co-localized marker pairs fall from a median *χ* of 9.6 × 10^3^ to 6.2 × 10^3^ *µ*m^2^, while opposed pairs rise from −8.1 × 10^3^ to −5.4 × 10^3^ *µ*m^2^; every one of the fifteen moves toward zero (Fig. 4A). The largest individual changes are *Cyp1a2* –*Cyp2a5* among the co-localized pairs (1.90×) and *Cps1* –*Cyp2a5* among the anti-localized ones (1.97×). If we rank instead by reproducibility across tissue cores the leading pairs are *Cyp1a2* –*Cyp2a5* (*Z*_*χ*_ = +2.03) and *Cyp1a2* –*Hal* (*Z*_*χ*_ = −1.48). At the level of individual mice, the coupling weakens in both CCl_4_ mice compared with both control mice for 11 of the 15 pairs, including every pair that involves *Cyp1a2* or *Cyp2a5*. This is the same architectural change found by the single-gene analysis of the previous section, now seen between genes. The two analyses agree: across tissue cores, the coupling of a marker pair follows the *χ* of its two genes.

The construction also assigns a zone to genes that carry no zonal annotation. Zonation markers differ in expression level along the porto-central axis and are not strictly present or absent, so a zone here means the side of the axis where expression is higher. A gene co-localized with a periportal marker is assigned to the periportal side, and a gene anti-localized with a periportal marker to the pericentral side Applying this criterion to every pair in which exactly one partner is an annotated marker places 1,266 previously unannotated genes on the zonation axis, 1,068 pericentral and 198 periportal (Fig. 4C; Supplemental Table S2). The imbalance reflects the composition of the two pair sets rather than the tissue. No gene receives votes for both zones, but the support varies: 582 genes are placed by a single marker pair, and 684 by two to six. Seven genes drawn from across both sets — *Apoa4, Etnppl, Acly* and *Gstp1* called periportal, *Abcb4, Cyp8b1* and *Lrrk1* called pericentral — were checked against published porto-central expression profiles from independent liver atlases (Yuan et al. 2025; Yakubovsky et al. 2026), and all seven agree with the assignment. The gene-pair correlation function can thus be used not only to detect the degradation of the zonation axis but to position unannotated genes along it.

### Relation to Moran’s *I*: the correlation function adds a length scale

Finally, we place the correlation-function quantities in the context of Moran’s *I* (Moran 1950), *a standard measure of spatial autocorrelation. In spatial transcriptomics Moran’s I* is normally computed on a nearest-neighbor graph and reported as a single number per gene, for example to detect spatially variable genes (Palla et al. 2022). To analyze the relation between 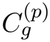 and Moran’s coefficient *I*, we computed *I* on the *k*-nearest-neighbor graph of hepatocytes (*k* = 6, each cell connected to its six closest cells, the default in Squidpy (Palla et al. 2022)), which in these tissues links cells less than about 30 *µ*m apart, in each of the seven control tissue cores (two mice), and compared it with the susceptibility *χ*_*g*_ of Eq (3) and the correlation length *ξ*_*g*_. The comparison is restricted to the 69 genes with an accepted exponential fit in all seven control cores and mean *R*^2^ *>* 0.7, so that *ξ*_*g*_ is meaningful for every gene considered. Across these genes Moran’s *I* follows the susceptibility closely (Spearman *ρ* = 0.89) but is unrelated to the correlation length (*ρ* = 0.18; Fig. 5A). The nearest-neighbor statistic thus reports how strongly the expression of neighboring cells is correlated, but carries little information on the distance over which that correlation persists. Since the susceptibility is only partly associated with the correlation length (*ρ* = 0.51), Moran’s I does not provide the same type of information that our method provides. Indeed, *χ*_*g*_ sums the correlation out to 100 *µ*m, so it accounts for both the strength and the range of the correlation. The two statistics are closely related. If Moran’s *I* is computed with a weight matrix that links every pair of cells separated by *s*_0_ to *R*_*χ*_, instead of the six nearest neighbors, it becomes the average of 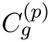 over this range weighted by the number of pairs, and in a tissue of uniform density it is approximately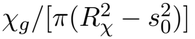. The susceptibility is therefore a distance-band Moran’s *I* in physical units; what the correlation function adds is the dependence on distance, and with it the correlation length.

**Figure 5.**
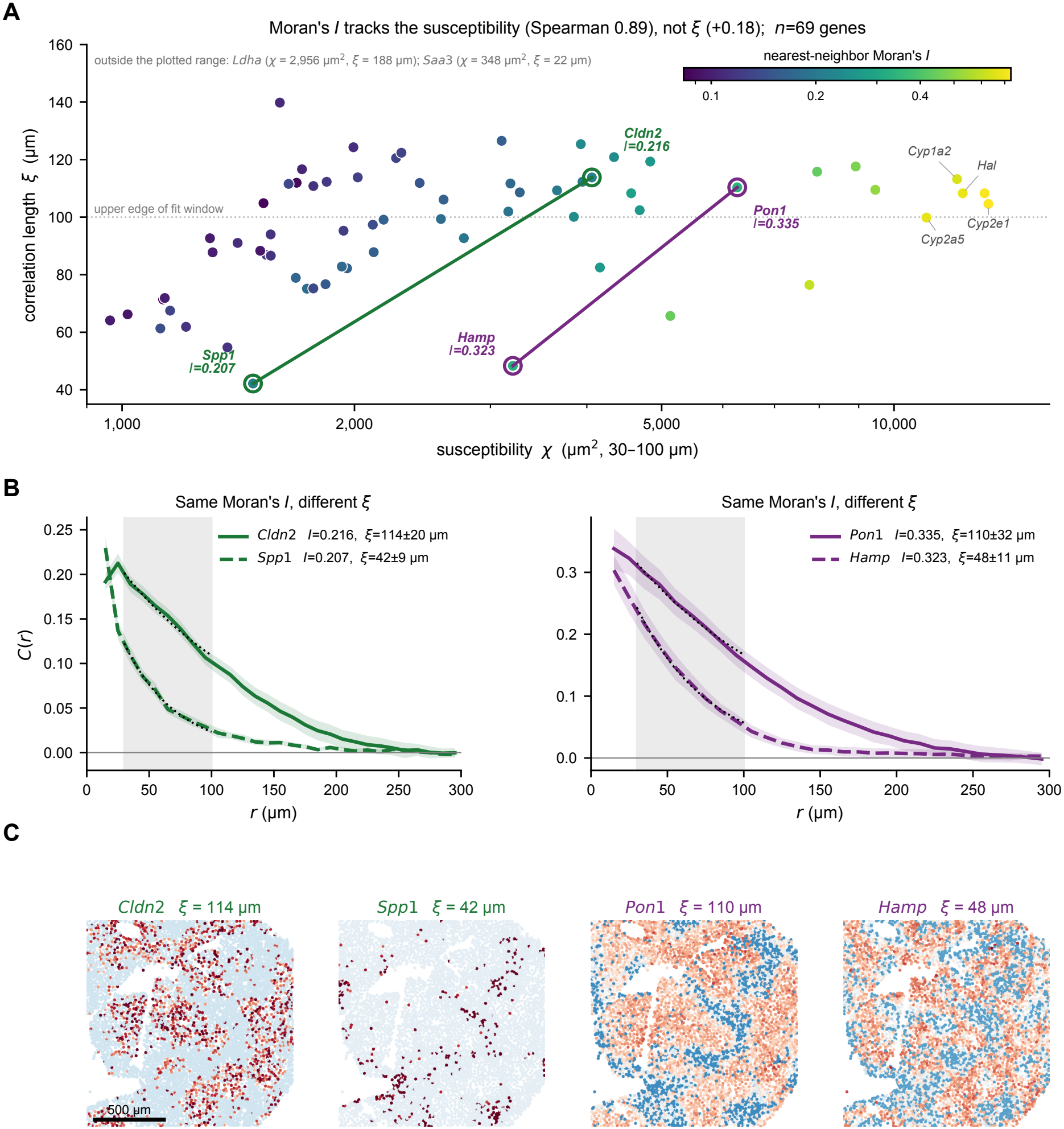
Moran’s *I* follows the susceptibility but not the correlation length. Mouse Xenium data, hepatocytes of the seven control tissue cores (two mice). (A) Susceptibility *χ*_*g*_ (Eq 3, 30–100 *µ*m) against the correlation length *ξ*_*g*_ for the 69 genes with an accepted exponential fit in every control core (mean *R*^2^ *>* 0.7); both are means over cores. Color gives Moran’s *I* computed on the *k*-nearest-neighbor graph (*k* = 6). Moran’s *I* increases with *χ*_*g*_ (Spearman *ρ* = 0.89) and is unrelated to *ξ*_*g*_ (*ρ* = 0.18). The dotted line marks the upper edge of the 30–100 *µ*m fit window; values of *ξ*_*g*_ above it are extrapolated. Circled points joined by a line are two pairs of genes with nearly equal Moran’s *I* and different *ξ*_*g*_; curated zonation markers are labeled in gray. (B) Correlation functions of the two pairs (mean over cores; shading, standard error). Dotted black lines show the exponential decay with the fitted *ξ*_*g*_ over the shaded 30–100 *µ*m window; values in the legend are mean ± s.d. of *ξ*_*g*_ across the seven control cores. (C) Expression of the four genes in one control core (`core_1`, mouse R1), standardized within the core; scale bar, 500 *µ*m.

Genes with the same Moran’s *I* can consequently be organized on different scales. For instance, *Cldn2* and *Spp1* have nearly equal values (*I* = 0.216 and 0.207) but correlation lengths of 114 ± 20 and 42 ± 9 *µ*m (mean ± s.d. across the seven control cores), and *Pon1* and *Hamp* (*I* = 0.335 and 0.323) have 110 ± 32 and 48 ± 11 *µ*m (Fig. 5B). Because *χ*_*g*_ accumulates the correlation up to 100 *µ*m, the longer-range gene of each pair also has two to three times the susceptibility (4,060 versus 1,478 *µ*m^2^, and 6,265 versus 3,208 *µ*m^2^), a difference that the nearest-neighbor value does not show. The difference is visible in the tissue: *Cldn2* and *Pon1* occupy broad territories, whereas *Hamp* varies on a finer scale and *Spp1* is confined to small isolated clusters (Fig. 5C). Note that the members of each pair coincide at the shortest separations, which is what the nearest-neighbor statistic measures, but already differ at 30–50 *µ*m. For 35 of the 69 genes *ξ*_*g*_ exceeds 100 *µ*m, the upper limit of the fit window, so the larger values are extrapolated (Methods); the comparison is between shorter and longer decay and not between precisely determined lengths. With these limits, the strength and the range of the correlation are two separate properties of a spatial expression pattern, and a single nearest-neighbor coefficient captures only the strength.

### Chronic human fibrosis causes a widespread loss of spatial correlation

We next applied the same analysis to a published multiplexed error-robust fluorescence in situ hybridization (MERFISH) dataset of healthy and fibrotic human liver (Watson et al. 2025), reanalyzed here without modification of the upstream processing. The contrast with the mouse cohort is strong. Of the 51 genes passing the filters, 50 (98%) show a correlation loss, with a median ratio of 4.9 (Fig. 6A) — against 49% and 1.42× in mouse. Chronic human fibrosis is therefore associated with a global contraction of spatial correlation across the measured panel, whereas repeated CCl_4_ injury in mouse produces a reorganization in which as many genes gain spatial correlation as lose it. Because the MERFISH panel was designed around genes that differ between hepatocyte zones in healthy liver (Methods), part of this contrast likely reflect the choice of genes.

**Figure 6.**
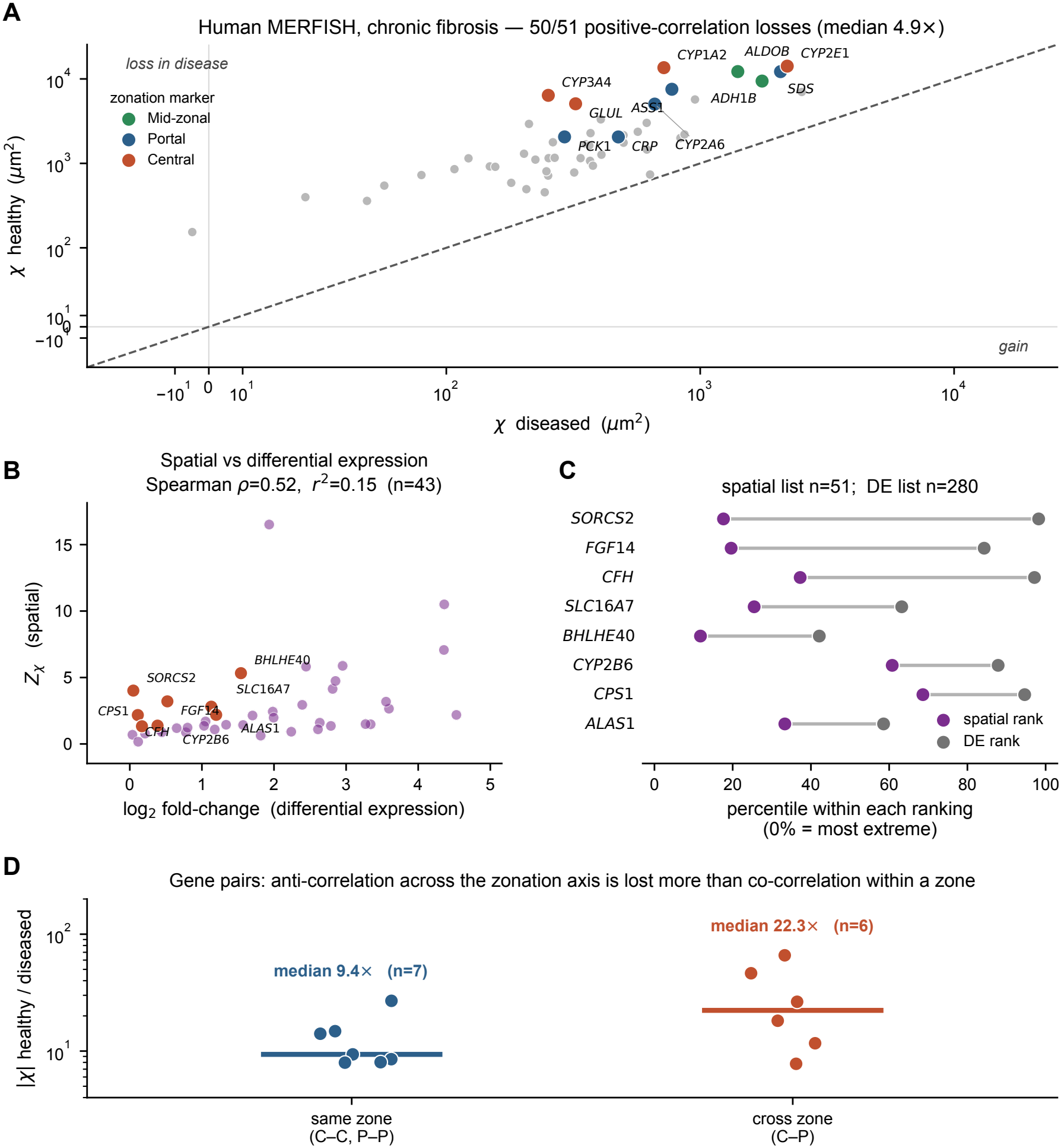
Chronic human fibrosis causes a widespread loss of spatial correlation. (A) Susceptibility *χ* per gene in healthy versus diseased tissue, computed and displayed as in Fig. 3A. 50 of 51 genes show a positive-correlation loss, with a median ratio of 4.9×. Curated zonation markers are colored by zone; pericentral, periportal and mid-zonal markers are all affected. (B) *Z*_*χ*_ against log_2_ fold-change. The two are moderately rank-associated but share little variance. (C) Position of the eight most discordant genes within each ranking, as in Fig. 3C. (D) Gene pairs built from the curated markers. Pairs spanning the pericentral–periportal axis lose more of their |*χ*| than pairs within a zone; each point is one pair and bars are medians. Of the pairs formed by the nine pericentral and periportal markers, only the 13 that pass the fit filters (Methods) are shown (7 within a zone, of which 6 are periportal, and 6 across zones); *GLUL* and *CRP* form no pair that passes.

Curated hepatocyte zonation markers are once again prominent and span the whole axis. Eleven of the 21 markers in the reference set (Watson et al. 2025) appear in the loss list: pericentral markers (*CYP3A4, CYP2E1, CYP1A2, GLUL*), periportal markers (*CYP2A6, SDS, ASS1, PCK1, CRP*) and the mid-zonal pair (*ALDOB, ADH1B*) all lose spatial correlation simultaneously, by factors ranging from 4.3 (*CRP*) to 25 (*CYP3A4*), with a median of 7.6. As in mouse, the relationship to differential expression is weak: the two rankings are moderately associated (*ρ* = 0.52, *p <* 0.001) but share little variance (*r*^2^ = 0.15), and individual genes are strongly discordant. For instance, *SORCS2* ranks in the top 18% spatially but the bottom 2% by DE, and *FGF14* and *CFH* behave likewise (Fig. 6B,C). Eight of the 51 genes in the human spatial list do not appear in the fold-change list at all. The highest-ranking genes are given in Table 3, and their spatial expression is shown in Supplemental Fig. S3.

**Table 3.**
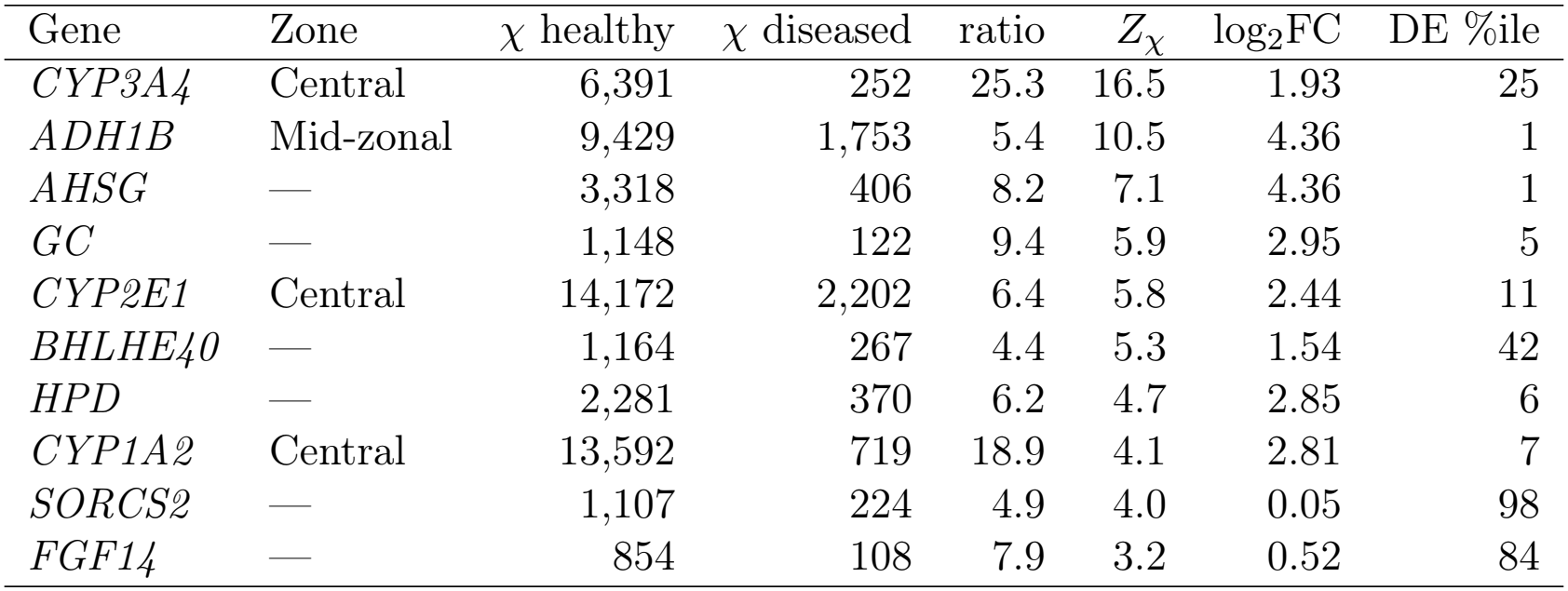
Human chronic fibrosis (MERFISH): genes with the largest loss of spatial correlation in fibrosis. The ten highest-ranking genes by *Z*_*χ*_. *χ* is the susceptibility (Eq 3) averaged over the samples of each condition; the ratio is *χ*_healthy_*/χ*_diseased_, so values *>* 1 indicate a loss of spatial correlation in disease. *Z*_*χ*_ (Eq 5) normalizes the difference by the between-sample variability. log_2_FC is the differential-expression fold change for the same cells, positive when the gene is more highly expressed in healthy tissue, and “DE %ile” is the gene’s percentile within the fold-change ranking (0% = largest fold-change); a high DE percentile therefore identifies a gene that reorganizes spatially while its mean abundance barely changes. “n.d.” marks genes whose *χ* is not positive in both conditions, for which a ratio is not meaningful. Across the whole panel, 50 of 51 genes show a positive-correlation loss (median 4.9×). Complete statistics for all genes are given in Supplemental Table S3.

| Gene | Zone | χ healthy | χ diseased | ratio | *Z*<sub>χ</sub> | log<sub>2</sub>FC | DE %ile |
|---|---|---|---|---|---|---|---|
| *CYP3A4* | Central | 6,391 | 252 | 25.3 | 16.5 | 1.93 | 25 |
| *ADH1B* | Mid-zonal | 9,429 | 1,753 | 5.4 | 10.5 | 4.36 | 1 |
| *AHSG* | — | 3,318 | 406 | 8.2 | 7.1 | 4.36 | 1 |
| *GC* | — | 1,148 | 122 | 9.4 | 5.9 | 2.95 | 5 |
| *CYP2E1* | Central | 14,172 | 2,202 | 6.4 | 5.8 | 2.44 | 11 |
| *BHLHE40* | — | 1,164 | 267 | 4.4 | 5.3 | 1.54 | 42 |
| *HPD* | — | 2,281 | 370 | 6.2 | 4.7 | 2.85 | 6 |
| *CYP1A2* | Central | 13,592 | 719 | 18.9 | 4.1 | 2.81 | 7 |
| *SORCS2* | — | 1,107 | 224 | 4.9 | 4.0 | 0.05 | 98 |
| *FGF14* | — | 854 | 108 | 7.9 | 3.2 | 0.52 | 84 |

The magnitude of this effect requires controls, because a change in *χ* could in principle arise from differences in sampling rather than in spatial organization. Neither the number of hepatocytes nor the variance normalization accounts for it. The fibrotic samples studied here in fact contain more hepatocytes than healthy ones (on average 65,500 versus 18,800, at about 1.7 times higher density), and 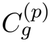 is a per-pair average, so cell number affects its precision but not its expectation: repeating the comparison with every sample subsampled to an identical 15,000 hepatocytes leaves the loss intact (median ratio 7.5 across the ten genes of Table 3, against 7.2 with all cells). Integrating the raw, unnormalized covariance instead of 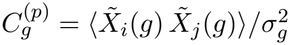 gives a similar loss (median ratio 7.9 for the same genes, and 5.2 against 4.9 across all 51 genes), so the normalization does not create the effect.

A third factor requires more care. The genes that lose spatial correlation are also detected in far fewer fibrotic hepatocytes (for example *CYP3A4* in 66% of healthy versus 19% of fibrotic hepatocytes), and sparser sampling depresses a measured correlation even when the underlying pattern is unchanged. The measured signal depends both on which cells have detectable expression of a gene and on the expression level in those cells. When the correlation function is restricted to cells with nonzero expression, six of seven zonation genes still lose spatial correlation, by factors of 2.6 to 4.9 (*CYP1A2* 4.9×, *CYP2E1* 4.7×, *ADH1B* 4.2×, *GLUL* 3.2×, *CYP3A4* 3.0×, *AHSG* 2.6×); only *GC* shows no residual change (0.9×), so its loss comes entirely from the fall in the fraction of cells with detectable expression.

Finally, the gene-pair analysis makes the architectural interpretation explicit. Among the curated markers, pairs drawn from the same zone lose a median of 9.4× of their |*χ*|, whereas pairs that are opposite along the pericentral–periportal axis lose 22.3× (Fig. 6D). The anti-correlation along the axis degrades more than the co-correlation within either pole. This contrast is confined to the strongly correlated pairs, which are the canonical zonation markers: split by sign alone across all 663 pairs, anti-correlated and co-correlated pairs lose comparably (6.3 versus 6.7×), and the asymmetry emerges only in the pairs with the highest |*χ*|.

## Discussion

We have introduced an approach to the analysis of spatial transcriptomics data that is inspired by correlation functions defined in statistical physics to describe liquids, magnets and other complex materials. For each gene, or pair of genes, the correlation function describes how the expression in one cell is related to the expression in other cells as a function of their distance. We condense each function into two numbers: (i) the correlation length *ξ*, the distance over which this relation decays, which measures how large the regions of coordinated expression are, and (ii) the susceptibility *χ*, which adds up the correlation over a fixed range of distances and so measures how strongly, overall, a cell shares its expression state with the cells around it. Both are in physical units, and *χ* does not depend on the quality of a fit, which makes it the more robust of the two in noisy tissue. The correlation length is the more intuitive of the two quantities but also the harder to estimate. In our data the exponential fit is limited by the noise of single-cell measurements and by the size of the tissue cores, which span only a few correlation lengths. Larger tissue sections, more biological replicates and platforms with higher detection efficiency should make *ξ* more precise, so that it can be used as a primary measure rather than alongside *χ*.

Several computational tools describe spatial organization in spatial transcriptomics data. Giotto (Dries et al. 2021) and Squidpy (Palla et al. 2022) focus on neighborhood enrichment, co-occurrence and cell–cell proximity. SpatialDE (Svensson et al. 2018), trendsceek (Edsgärd et al. 2018), SPARK (Sun et al. 2020) and SPARK-X (Zhu et al. 2021) test whether a gene is spatially variable, and SpatialDE also estimates a length scale for each gene. CELINA (Shang et al. 2025) extends this test to genes that vary within a single cell type, as we did here by restricting the analysis to hepatocytes. Statistics from the geographical sciences, such as Moran’s *I*, Geary’s *C* and the Getis–Ord *G* statistic, have been proposed as a general framework for these data (Zormpas et al. 2023). The closest of these to our approach is Moran’s *I* (Moran 1950), *often used to detect spatially variable genes. It is usually computed on the graph of nearest neighbors, and in that form it follows the susceptibility closely but carries no information on the correlation length (Fig. 5): genes with the same Moran’s I* can be organized over very different distances. The correlation function makes this distance explicit, because it is evaluated at every separation rather than at one choice of neighborhood.

Two recent methods also measure spatial correlations in gene expression. spCorr (Jiang et al. 2026) maps how the correlation between two genes measured in the same spot or cell varies across a tissue, and the network pair correlation function (netPCF) (Moore et al. 2026) counts how often cells with given expression levels are found at distance *r*, measuring distance along a network of neighboring cells to correct for irregular tissue geometry. Our functions instead relate the continuous value of the expression of a gene in one cell to that of the same or a second gene in other cells, and give two numbers that can be computed for every gene and gene pair of a panel and compared directly between conditions, as we did here for more than 250,000 gene pairs. The approaches are complementary: spCorr could show where in the lobule the coupling between zonation markers is lost, and network distances could account for vessels and septa, while *χ* and *ξ* summarize, for each gene or gene pair and over the whole sample, how much spatial coordination is lost and over what distance.

The liver is a natural test case for these quantities, because few tissues are as spatially organized. Hepatocytes are arranged along the axis that runs from the portal triads, where branches of the portal vein, hepatic artery and bile duct run together, to the central vein of each lobule, and their gene expression changes gradually along this axis (Halpern et al. 2017). In the mouse this axis is about 210 *µ*m long (Ruijter et al. 2004), so the distances summed in *χ*, up to 100 *µ*m, cover about half of it. A gene that follows the zonation pattern therefore has a large susceptibility and a correlation length of about 100 *µ*m in healthy tissue, and any disease that disturbs zonation should show up as a change in these two numbers.

CCl_4_ injury is a clear example. CCl_4_ is activated by cytochrome P450 2E1 (CYP2E1), which is expressed mainly in pericentral hepatocytes (Albadry et al. 2024), and its metabolites cause lipid peroxidation and hepatocyte damage concentrated in the pericentral region of the lobule (Weber et al. 2003), where fibrosis then begins (Geerts et al. 1991). Consistent with this, the correlation length of several cytochrome P450 genes is shorter in CCl_4_-treated mice (for example *Cyp2a5*, 100 to 72 *µ*m, and *Cyp2e1*, 105 to 74 *µ*m; Figs. 1 and 2), whereas that of the periportal and midzonal genes *Arg1, Cps1, Etnppl* and *Hamp* is unchanged. Among the genes placed on the zonation axis by the gene-pair analysis, *ξ* shortens for those assigned to the pericentral side (median ratio 1.14, 100 genes) but not for those assigned to the periportal side (0.84, 50 genes). The contraction of correlated domains is therefore concentrated in pericentral hepatocytes, where the toxic effect of CCl_4_ is expected, although the susceptibility of periportal markers also decreases and *Hal* is an exception among them. CCl_4_ also causes oxidative stress (Knockaert et al. 2012), which activates the transcription factor Nrf2, the main regulator of the antioxidant response (Nguyen et al. 2009). *Cyp2a5*, the gene whose correlation length contracts most clearly, is a target of Nrf2 (Lämsä et al. 2010), whereas *Etnppl*, whose correlation length is unchanged, encodes an enzyme of phospholipid metabolism (A Elmihi et al. 2024). Changes in spatial correlation may thus separate genes that respond to an external stress from genes whose organization is preserved. These changes are only weakly related to changes in expression level (Fig. 3): genes such as *Cyp2c29* and *Pon1* lose spatial correlation in both CCl_4_-treated mice while their average expression does not consistently fall, so a ranking by fold-change would not bring them forward. *Cyp2c29* in particular protects against liver inflammation after CCl_4_ injury (Wang et al. 2020), so a change in its spatial organization may be relevant to the injury response.

The gene-pair correlations add a second type of information: the sign of 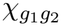 tells whether two genes are co-localized or anti-localized. Pairs of known markers from the same zone have positive coupling and pairs from opposite zones have negative coupling, in every tissue core (Fig. 4). A gene that is not a known marker can therefore be placed on the zonation axis from the sign of its coupling with the known markers, and we placed 1,266 such genes in this way. The calls agreed with published zonation profiles for all seven genes we checked.

In human fibrosis the loss of spatial correlation is much larger and more uniform: 50 of the 51 genes analyzed lose susceptibility, with a median ratio of 4.9, against 49% of genes and a median ratio of 1.42 in mouse. The controls in Results show that this loss is not explained by the number of hepatocytes, the variance normalization or the fall in detection in fibrotic tissue. The two gene panels overlap only partially, because the MERFISH panel has 317 hepatocyte-focused genes and the Xenium panel is genome-scale, so after the quality filters only six orthologous genes are measured in both. Within this small set, all six genes lose susceptibility in both species, but by very different amounts: the median susceptibility ratio is 6.1 in human against 1.7 in mouse. *CYP1A2* falls by a factor of 18.9 in human but only 1.9 in mouse, and *CYP2E1* by 6.4 against 1.3.

Several differences between the two cohorts could contribute to this contrast. The first is the choice of genes: the MERFISH panel was selected among genes that differ between hepatocyte clusters in healthy liver (Methods), so it is enriched for strongly zonated genes, which are also the genes with the largest susceptibility in healthy tissue. Part of the difference in the fraction of genes that lose spatial correlation, 98% in human and 49% in mouse, may therefore reflect the panel rather than the injury. A second possibility is the size of the cells. Human hepatocytes are larger than mouse hepatocytes, but only by a factor of about 2.2 in volume, or 1.3 in diameter (20–30 against 15–20 *µ*m) (Hammad et al. 2014; Jiang et al. 2020; Rohr et al. 1976), and most mouse hepatocytes are binucleated whereas most human hepatocytes are mononucleated (Celton-Morizur et al. 2009; Duncan et al. 2010; Gentric et al. 2015). Because each susceptibility ratio compares two conditions of the same species, with the same distance cutoffs, cell size alone is unlikely to explain the difference. The most likely remaining factor is the stage of the injury. Three weeks of CCl_4_ treatment produces an early stage of fibrosis that regresses after the treatment is stopped, whereas the human samples come from established fibrosis of stages F2 to F4, after long-standing injury, when regression is slower and less complete (Iredale 2007; Kisseleva and Brenner 2021). The mouse tissue is therefore closer to its healthy organization, and its smaller loss of spatial correlation is consistent with this difference in stage. The same pattern holds for gene pairs. In both species the anti-correlated pairs, which link genes of opposite zones, weaken more often than the co-correlated ones, but at very different amplitudes. In human fibrosis 97% of all pairs weaken, with a median |*χ*| ratio of 6.6, whereas in mouse 65% of the anti-correlated and 46% of the co-correlated pairs weaken, with median ratios of 1.30 and 0.95. The two injuries thus agree in direction but differ by about an order of magnitude in effect: in the mouse the zonation axis is weakened, while in human fibrosis, at a more advanced stage, it is largely lost. If this difference reflects the stage of the disease, *χ* and *ξ* could be used to grade how far zonation has been lost, which could be tested by following the same model through longer treatment and through recovery. Note that the mouse comparison rests here on two animals per condition, so its effects are reported as consistent across mice rather than tested statistically, and both cohorts are single time points. Following the same tissue through the course of injury would show whether the loss of correlation precedes, accompanies or follows the fibrotic remodeling.

Correlation functions have been used to characterize spatial organization of liquids and magnets using a few measurable quantities. Applied to spatial transcriptomics, they do the same for tissues: they measure how strongly and over what distance cells coordinate their gene expression, and they show how this coordination is lost in disease even when the average expression of a gene barely changes. Because they need only the position and expression of each cell, they can be applied to other tissues in which disease changes spatial organization, such as fibrosis of the heart, kidney or lung.

## Methods

### Animals and CCl_4_ treatment

C57BL/6J mice were bred and housed at the CEBEGA-USC animal facility. All procedures were authorized by the Animal Ethics Committee of the University of Santiago de Compostela and the Xunta de Galicia and were conducted in accordance with applicable national and international regulations for the care and use of laboratory animals. Animals were housed in groups of up to five per cage with wood-chip bedding, under controlled temperature (20–22 ^*°*^C) and relative humidity (45 ± 10%), with a 12 h light/12 h dark cycle (lights on from 08:00 to 20:00). Water and standard rodent chow were provided ad libitum. The maintenance diets used were Teklad Global 14% Protein Rodent Maintenance Diet 2914 or Teklad Global 18% Protein Rodent Diet 2918 (Envigo).

To induce liver injury, mice received intraperitoneal injections of carbon tetrachloride (CCl_4_; 06-3545, Strem Chemicals) twice weekly for three consecutive weeks. Both treatment and control mice received intraperitoneal injections. The treatment group received a dose of 200 *µ*L/kg body weight of CCl_4_ diluted 1:50 in corn oil (C8267, Sigma-Aldrich), whereas the control group received an equivalent volume of corn oil alone. Animals were fasted for 16 h before sample collection. Prior to blood collection, mice were anesthetized with 4% isoflurane, and blood was obtained from the retro-orbital sinus using heparin-coated glass capillaries (2901000, Marienfeld). After collection, the liver was removed and fixed overnight at 4 ^*°*^C in 4% paraformaldehyde prepared in PBS (sc-281692, Santa Cruz Biotechnology). The tissue was subsequently washed three times with calcium- and magnesium-free DPBS (14190-094, Gibco).

### Xenium spatial transcriptomics of mouse liver

Spatially resolved single-cell transcriptomes of mouse liver were generated using the Xenium Prime In Situ Gene Expression workflow for FFPE tissue (10X Genomics). We used the Xenium Prime 5K Mouse Pan Tissue & Pathways Panel (10X Genomics, 1000725, RevE, 5,006 genes) supplemented with a 50-gene custom add-on panel (design ID V9FQHP), giving 5,056 targeted genes in total. Livers were obtained from male C57BL/6J mice aged 10 weeks that received intraperitoneal injections twice weekly for three consecutive weeks of either CCl_4_ (200 *µ*L/kg body weight, diluted 1:50 in corn oil; *n* = 2) or an equivalent volume of corn oil alone (controls; *n* = 2), as described above. Livers were fixed overnight at 4 ^*°*^C in 4% paraformaldehyde in PBS, washed three times in calcium- and magnesium-free DPBS and paraffin-embedded. Cores of 2 mm diameter from the paraffin blocks were assembled into a tissue microarray (TMA). The TMA was sectioned at 5 *µ*m, and sections were placed in the sample area of a Xenium slide following the Tissue Preparation Guide (10X Genomics, CG000578). Slides were dried for 3 h at 42 ^*°*^C and then placed in a desiccator at room temperature overnight. Slides were fixed and permeabilized according to the Xenium In Situ Gene Expression Fixation and Permeabilization protocol (10X Genomics, CG000580, RevE). Probe hybridization, ligation, rolling circle amplification and cell segmentation staining followed the manufacturer’s instructions in Xenium In Situ Gene Expression with optional Cell Segmentation Staining (10X Genomics, CG000760, RevA); the custom add-on panel was added at the priming hybridization and probe hybridization steps. Slides were imaged on the Xenium Analyzer (10X Genomics, CG000584, RevF; instrument software v4.0.1.4) with a pixel size of 0.2125 *µ*m and a z-step of 3 *µ*m.

Raw data were processed with Xenium Onboard Analysis v4.0.1. This step performed transcript decoding and multimodal cell segmentation based on the Xenium Multi-Tissue Stain (ATP1A1/CD45/E-Cadherin boundary stain, 18S interior stain and DAPI nuclear stain, with a 5 *µ*m nuclear expansion for cells lacking a boundary signal). Only transcripts with a quality score

≥ 20 were assigned to cells. The run yielded 422,157 segmented cells across the whole slide. Liver cells were selected with a manually drawn polygon on the cell centroid coordinates, and cells were assigned to individual TMA cores by DBSCAN clustering of centroids (eps = 200 *µ*m, minimum 5 cells). Cores with ≥ 500 cells were matched to their samples, and detached tissue fragments were excluded. The final dataset comprised six samples (four control, two CCl_4_) corresponding to six paraffin blocks from four mice (two per condition; blocks from the same mouse are denoted .1/.2), represented by 11 TMA cores. Cells with fewer than 10 transcripts or fewer than 5 detected genes were removed, as were genes detected in fewer than 10 cells. Raw counts were normalized to 10,000 counts per cell and log-transformed (log1p). Using raw counts, we selected 2,000 highly variable genes (Seurat v3 method) and computed 50 principal components. A neighborhood graph was built from the first 30 principal components (15 neighbors) and used for UMAP embedding and Leiden clustering (resolution 0.5). Clusters were annotated into hepatocytes, endothelial cells, stellate cells/fibroblasts, Kupffer cells/macrophages and cholangiocytes using canonical marker genes (*Apob, Hamp, Cyp2e1, Arg1, Hal, Cps1, Kdr, Pecam1, Lyve1, Dcn, Col1a1, Csf1r, Cd163, Epcam, Spp1*), and two clusters with very low transcript counts were labeled as low quality. Analyses were performed in Python using Scanpy (v1.10.2).

### Tissue cores used in the correlation analysis

The correlation analysis uses the hepatocytes of ten of the eleven tissue cores: seven from the two control mice (R1: three cores, from blocks R1.1 and R1.2; R3: four cores, from blocks R3.1 and R3.2) and three from the two CCl_4_-treated mice (R2: two cores, from block R2.2; R4: one core), 74,800 hepatocytes in total. The eleventh core, the second core of mouse R4, was excluded by hand because it contained only 584 hepatocytes, too few for a reliable estimate of the correlation function, and had very low transcript detection (median 66 counts per cell). Mouse R4 as a whole has much lower transcript detection than the other mice (median of about 100 counts per cell in its included core, against 530 to 950 in the other cores). Note that cores from the same mouse, and in particular from the same block, are not equivalent to independent biological replicates.

### MERFISH dataset

Spatially resolved single-cell transcriptomes were obtained by multiplexed error-robust fluorescence in situ hybridization (MERFISH) of human liver tissue, originally generated by Watson et al. (2025). The gene panel comprised 317 genes with an intentional focus on hepatocyte-expressed genes: they were selected among the genes differentially expressed between hepatocyte clusters in single-cell, single-nucleus and spatial RNA sequencing of healthy human liver (Andrews et al. 2022), supplemented with common markers of non-parenchymal liver cells (MacParland et al. 2018). The panel was therefore designed from healthy tissue, and not selected for fibrosis-related genes. Eight liver samples were included in our analysis: three healthy samples and five fibrotic samples. The original study performed cell-type annotation using two complementary integration strategies. In the healthy-only analysis, three hepatocyte populations reflecting canonical zonation were identified: Hep 1, Hep 2, and Hep 3. In the joint healthy-and-diseased analysis, hepatocytes were organized into two healthy-associated populations (Portal Hep and Central Hep) and two fibrosis-associated populations (Fibrotic Hep 1 and Fibrotic Hep 2). We used the healthy-only labels for analyses restricted to healthy samples and the joint-analysis labels for diseased samples. For each sample, spatial coordinates correspond to cell centroids in micron units. The publicly released expression matrices were already normalized by total transcript counts per cell, scaled to a common library size, and log-transformed with a pseudocount. Prior to analysis, low-quality cells with extremely low transcript counts or missing cell-type annotations were removed. The MERFISH data were obtained from Dryad (https://doi.org/10.5061/dryad.37pvmcvsg).

### Computing correlation functions, susceptibility and correlation lengths

All symbols are listed in Table 4. For each sample, the input is a cell-by-gene expression matrix *X ∈* ℝ^*N ×G*^ and a set of two-dimensional cell centroid coordinates {**r**_*i*_}. Pairwise Euclidean distances are computed using a KDTree nearest-neighbor search (`sklearn.neighbors`). Cell pairs are assigned to radial distance bins of width Δ*s* = 5 *µ*m, defining the sets *D*_*p*_ used in the correlation functions of Eqs (1) and (4). Gene-expression vectors are mean-centered across cells, and 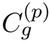 and 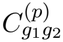 are then evaluated bin by bin. Throughout this work both the susceptibility of Eq (3) and the correlation length are evaluated over the same interval, *s*_0_ = 30 *µ*m to *R*_*χ*_ = 100 *µ*m, so that *ξ* and *χ* characterize the same range of separations. The upper cutoff is constrained from both sides. Explicitly, there is a sum rule for the correlation function which reads 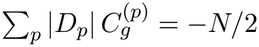 for a single gene, so *C*^(*p*)^ must turn negative at large separation whatever the underlying biology; in these tissues it crosses zero near 250 *µ*m. A cutoff of 100 *µ*m therefore stays well inside the positive, densely sampled short-range regime, where *χ* is dominated by genuine correlated structure rather than by the noisy tail or the sum-rule-induced negative region, and it corresponds physically to a few cell diameters, the scale of local tissue neighborhoods. Applying one fixed value to all genes, gene pairs and conditions, rather than a per-gene value tied to *ξ*, keeps susceptibilities on a common spatial scale; the ranking is in any case stable within this regime: with the lower cutoff between 20 and 50 *µ*m and the upper cutoff between 60 and 120 *µ*m, the rank correlation of *Z*_*χ*_ with the reported ranking is at least 0.96. Correlation lengths are extracted by fitting the decay of *C*^(*p*)^ with distance to an exponential model, using nonlinear least-squares optimization (`scipy.optimize.curve_fit`). We use the name susceptibility for *χ* by analogy with statistical physics, where, in thermal equilibrium, the integral of the correlation function is proportional to the response of the system to a uniform external field through the fluctuation–dissipation theorem (Chaikin and Lubensky 1995). A living tissue is not in thermal equilibrium and this relation does not apply, so here *χ* is defined only as the correlation summed over a range of distances and does not measure a response to an external perturbation.

**Table 4.**
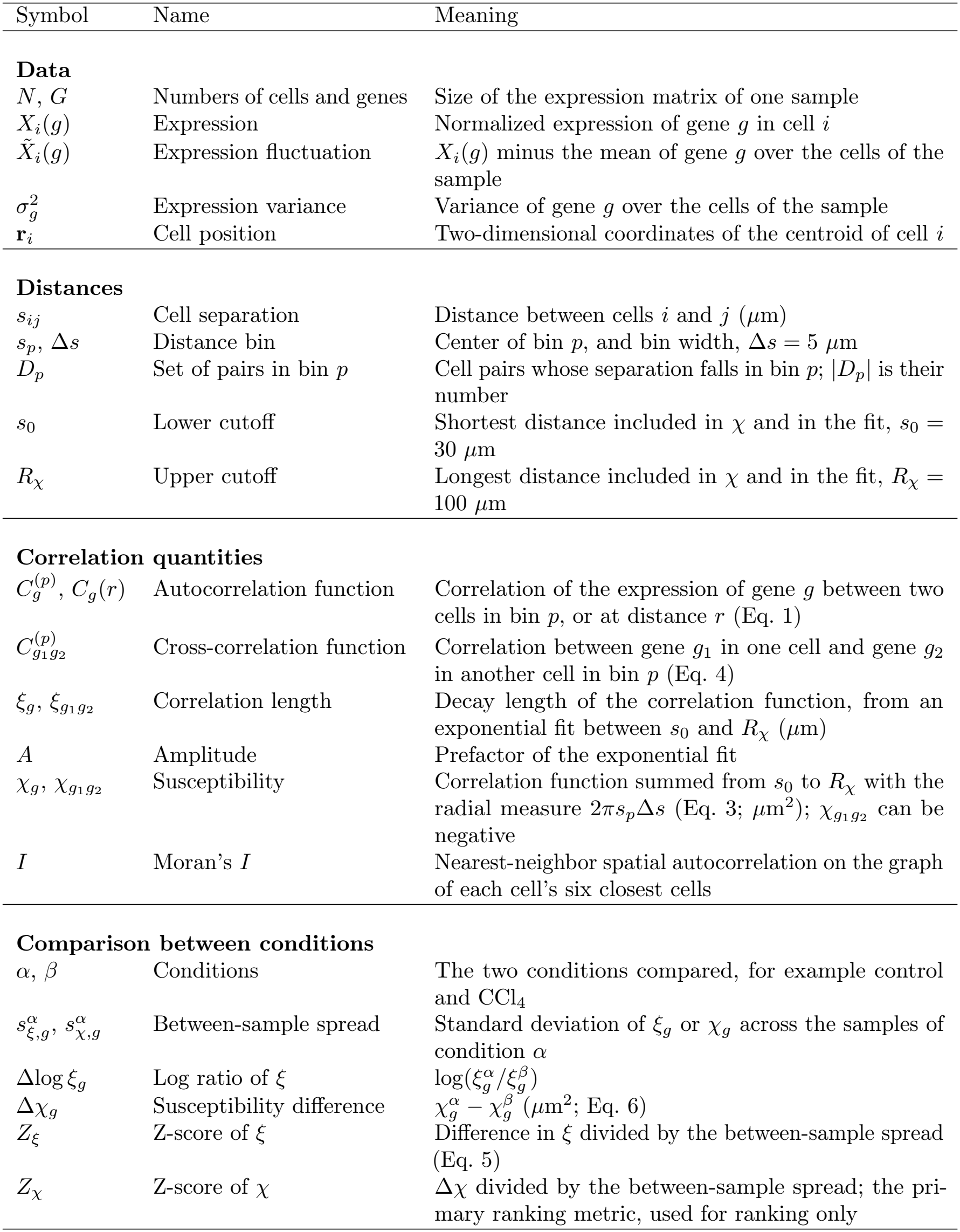
Symbols used in this work. Quantities without units are dimensionles.

| Symbol | Name | Meaning |
|---|---|---|
| **Data** |  |  |
| *N*, *G* | Numbers of cells and genes | Size of the expression matrix of one sample |
| *X*<sub>*i*</sub>(*g*) | Expression | Normalized expression of gene *g* in cell *i* |
| $\tilde{X}_i(g)$ | Expression fluctuation | *X*<sub>*i*</sub>(*g*) minus the mean of gene *g* over the cells of the sample |
| $\sigma_g^2$ | Expression variance | Variance of gene *g* over the cells of the sample |
| **r**<sub>*i*</sub> | Cell position | Two-dimensional coordinates of the centroid of cell *i* |
| **Distances** |  |  |
| *s*<sub>*ij*</sub> | Cell separation | Distance between cells *i* and *j* (*μ*m) |
| *s*<sub>*p*</sub>, Δ*s* | Distance bin | Center of bin *p*, and bin width, Δ*s* = 5 *μ*m |
| *D*<sub>*p*</sub> | Set of pairs in bin *p* | Cell pairs whose separation falls in bin *p*; \|*D*<sub>*p*</sub>\| is their number |
| *s*<sub>0</sub> | Lower cutoff | Shortest distance included in *χ* and in the fit, *s*<sub>0</sub> = 30 *μ*m |
| *R*<sub>*χ*</sub> | Upper cutoff | Longest distance included in *χ* and in the fit, *R*<sub>*χ*</sub> = 100 *μ*m |
| **Correlation quantities** |  |  |
| $C_g^{(p)}$, *C*<sub>*g*</sub>(*r*) | Autocorrelation function | Correlation of the expression of gene *g* between two cells in bin *p*, or at distance *r* (Eq. 1) |
| $C_{g_1g_2}^{(p)}$ | Cross-correlation function | Correlation between gene *g*<sub>1</sub> in one cell and gene *g*<sub>2</sub> in another cell in bin *p* (Eq. 4) |
| $\xi_g$, $\xi_{g_1g_2}$ | Correlation length | Decay length of the correlation function, from an exponential fit between *s*<sub>0</sub> and *R*<sub>*χ*</sub> (*μ*m) |
| *A* | Amplitude | Prefactor of the exponential fit |
| $\chi_g$, $\chi_{g_1g_2}$ | Susceptibility | Correlation function summed from *s*<sub>0</sub> to *R*<sub>*χ*</sub> with the radial measure 2*πs*<sub>*p*</sub>Δ*s* (Eq. 3; *μ*m<sup>2</sup>); $\chi_{g_1g_2}$ can be negative |
| *I* | Moran's *I* | Nearest-neighbor spatial autocorrelation on the graph of each cell's six closest cells |
| **Comparison between conditions** |  |  |
| *α*, *β* | Conditions | The two conditions compared, for example control and CCl<sub>4</sub> |
| $s^{\alpha}_{\xi,g}$, $s^{\alpha}_{\chi,g}$ | Between-sample spread | Standard deviation of *ξ*<sub>*g*</sub> or *χ*<sub>*g*</sub> across the samples of condition *α* |
| Δlog *ξ*<sub>*g*</sub> | Log ratio of *ξ* | $\log(\xi_g^{\alpha}/\xi_g^{\beta})$ |
| Δ*χ*<sub>*g*</sub> | Susceptibility difference | $\chi_g^{\alpha} - \chi_g^{\beta}$ (*μ*m<sup>2</sup>; Eq. 6) |
| *Z*<sub>*ξ*</sub> | Z-score of *ξ* | Difference in *ξ* divided by the between-sample spread (Eq. 5) |
| *Z*<sub>*χ*</sub> | Z-score of *χ* | Δ*χ* divided by the between-sample spread; the pri-mary ranking metric, used for ranking only |

### Fit quality and estimation of uncertainties

Fits are accepted only when *ξ >* 0, *ξ <* 5 *s*_max_ where *s*_max_ is the largest pair separation retained, and *R*^2^ *>* 0.1, with *R*^2^ the coefficient of determination. The exponential model here is used to provide an effective decay scale over the fit window rather than an exact description of the *C*^(*p*)^ functions. Even for genes with *R*^2^ *>* 0.98, the fitted *ξ* depends on the choice of fit window, varying by up to 20% across reasonable choices. Ratios of correlation lengths between two conditions measured with the same window are considerably more stable, varying by only a few percent. This is the rationale for the ranking metrics described below, which are built on the log ratio Δlog *ξ*_*g*_ and on the fit-independent susceptibility difference Δ*χ*_*g*_ rather than on the absolute values of *ξ*. Uncertainties on *ξ* are estimated as the standard deviation of the per-sample fitted values across samples (tissue cores for the Xenium dataset, which come from two mice per condition, and different samples for the MERFISH dataset). Only genes and pairs for which the *ξ* exponential fit is above the *R*^2^ cutoff for at least three samples are used in the calculation of the mean and standard deviation.

### Ranking genes and gene pairs by the change between conditions

To rank genes (or gene pairs) by the change in spatial organization between two conditions *α* and *β* (healthy and diseased hepatocytes) we use three complementary metrics. For the correlation length, the z-score is

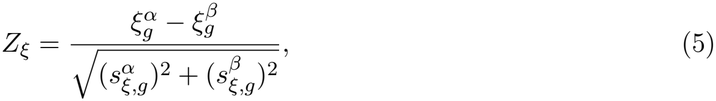

where 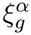 is the mean of *ξ*_*g*_ over the samples of condition *α* and 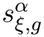 is its standard deviation across these samples (and likewise for *β*). Because the denominator is the sample-to-sample spread, *Z*_*ξ*_ measures reproducibility across samples and it down-weights genes whose correlation length varies substantially between samples. For the Xenium data the samples are tissue cores (seven control and three CCl_4_ cores from two mice per condition), which are not independent biological replicates, and for MERFISH they are *n* = 3 control and *n* = 5 fibrotic samples; in both cases *s*_*ξ,g*_ is imprecisely estimated, so *Z*_*ξ*_ should be read as a ranking statistic rather than as a calibrated test statistic: we use it to order genes and apply a permissive threshold (|*Z*_*ξ*_| *≳* 1). For reporting effect magnitude in a scale-invariant way, analogous to log fold-change in differential expression analysis, we use the log ratio 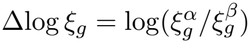. As a model-free alternative that does not rely on the exponential fit, we use the susceptibility difference, evaluated over the same range as Eq (3),

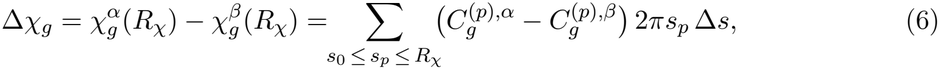

with the corresponding z-score 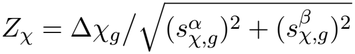, where 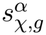 is the standard deviation of *χ*_*g*_ across the samples of condition *α. Z*_*χ*_ is the primary ranking metric used throughout this work.

The same metrics extend unchanged to the gene-pair statistics, giving 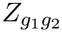, Δlog 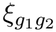 and 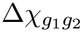. Because 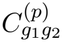 may be negative for genes expressed in opposite zones, gene pairs are ranked by 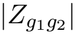 so that loss of positive and of negative coupling are treated symmetrically, and given the *∼ G*(*G*−1)*/*2 comparisons involved we report a fixed number of top-ranking pairs.

## Software availability

Code implementing the correlation functions and the estimators of *ξ* and *χ* is available at https://github.com/Krrsshh/ST-CorrelationFns.

## Data access

The mouse liver Xenium data generated in this study (control and CCl_4_) have been deposited in the BioImage Archive under accession S-BIAD4317 (doi:10.6019/S-BIAD4317; https://www.ebi.ac.uk/biostudies/bioimages/studies/S-BIAD4317) and will be available upon publication. The deposit includes transcript coordinates with quality scores, cell × gene count matrices, cell metadata, cell and nucleus boundary polygons, segmentation masks, morphology images cropped to each sample, and the probe panel. The same accession contains the processed and annotated single-cell object (120,256 cells, 5,056 genes, raw counts and normalized expression, main cell types), formatted according to the CELLxGENE schema and validated with the CELLxGENE validator. The processed object deposited in the BioImage Archive (S-BIAD4317) contains the 120,256 cells assigned to the six tissue blocks and the complete 5,056-gene panel. The analysis object comprised 123,853 cells and 5,046 genes after quality control; it additionally includes 3,597 cells from detached tissue fragments not assigned to any sample, and the 10 genes detected in fewer than 10 cells are included in the deposited raw counts but flagged as filtered. Per-gene results for both cohorts are provided as Supplemental Tables S1 and S3, and the zone assignments of Fig. 4C as Supplemental Table S2.

## Competing interest statement

The authors declare that they have no conflict of interest.

## Acknowledgments

We acknowledge support from the NIH/NIGMS under R35GM149261. This publication does not represent official views of the NIH. J.P. acknowledges support from Taighde Éireann - Research Ireland under the Government of Ireland Postdoctoral Fellowship Programme GOIPD/2025/1111. D.B. was supported by NIH/NIGMS 5T32GM154667-01. S.B. was supported in part by Ag-BioResearch at Michigan State University with funding from the Hatch Act capacity funding program (Accession Number 1018334). This research was supported by the Spanish Ministry of Science and Innovation-National Research Agency grants RED2022-134485-T (to M. V. R. and A. R. V.); RYC21-034611-I and CNS2022-135973 (to A. R. V.); the European Social Fund Plus (RYC21-034611-I); the European Union NextGenerationEU/Plan de Recuperación, Transforma-ción y Resiliencia (PRTR) (CNS2022-135973)(to A. R. V.); and the University of Málaga (Plan Propio, ayuda G) (to A. R. V.).

## Author contributions

C.P., S.B., and A.R.V. jointly supervised the work. A. R. V. and C.K. designed and supervised the Xenium experiment. A. R. V. discussed the biological aspects. C.P. and J.P. developed the correlation-function framework. K.C.G. implemented the correlation functions and carried out the analysis. D.B. analyzed the data and discussed biological aspects. J.C.M., A.C.D. and M.V.R. performed the CCl_4_ treatment and collected the mouse livers, and M.V.R. supervised the animal work. C.K. and J.C.F. processed the Xenium data, and J.C.F. annotated them and deposited them in public repositories. C.P. wrote the manuscript with input from all authors. We acknowledge the use of AI tools (Claude, Opus 5.5) to assist editing parts of the manuscript text and figures. The manuscript was reviewed by all the authors, who take full responsibility for its content.

## Notes

### Competing Interest Statement

The authors have declared no competing interest.

